# Trichostatin A Treatment Has Little Impact on Nuclear compartments in Cells Depleted of H3K9me, H3K27me3, and uH2A

**DOI:** 10.1101/2025.04.26.646718

**Authors:** Kei Fukuda, Chikako Shimura, Yoichi Shinkai

## Abstract

In the mammalian nucleus, heterochromatin is segregated from transcriptionally active euchromatic regions (A compartments), forming large, condensed, and inactive nuclear compartments (B compartments). However, the mechanisms underlying its spatial organization remain incompletely understood. We previously demonstrated that simultaneous depletion of H3K9 methylation, H3K27me3, and H2A monoubiquitination in immortalized mouse embryonic fibroblasts (iMEFs) leads to heterochromatin weakening and alterations in nuclear compartments. However, the overall pattern of A/B compartments remains largely preserved even under these conditions, suggesting the involvement of unknown factors. Since histone deacetylation is a key factor in transcriptional repression, we investigated the impact of histone deacetylase (HDAC) inhibition on nuclear compartments by treating cells depleted of H3K9 methylation, H3K27me3, and H2A K119 monoubiquitylation (H2AK119ub/uH2A) with the HDAC inhibitor Trichostatin A (TSA). We performed Hi-C analysis to assess the nuclear compartment changes. Our results showed that, although TSA treatment globally increased H3K27ac levels, the difference in H3K27ac enrichment between A/B compartments remained evident, and nuclear compartments were minimally affected. These findings suggest that factors other than HDACs maintain nuclear compartmentalization when repressive chromatin modifications are depleted.

## Description

Repressive chromatin modifications condense chromatin through their reader proteins (Bell et al. 2023). The representative repressive chromatin modifications in mammals—H3K9 methylation, H3K27me3, and uH2A—are mediated by SETDB1/SUV39H1/SUV39H2/EHMT1/EHMT2, EZH1/EZH2, and RING1A/RING1B, respectively. We generated cells deficient in all H3K9 methyltransferases and subsequently established 6KO cells lacking *Ring1b* (Fig. 1A) (Matsui et al. 2010; Kato et al. 2018; Fukuda et al. 2021; Fukuda et al. 2023). In these cells, simultaneous knockdown of *Ring1a* and treatment with the Ezh1/Ezh2 enzymatic inhibitor DS3201 successfully depleted all three repressive modifications (H3K9 methylation, H3K27me3, and uH2A) (Fukuda et al. 2025). However, even under these conditions, nuclear compartments were largely maintained (Fukuda et al. 2025) suggesting that other factors may be involved in the regulation of nuclear compartmentalization.

**Figure 1.**
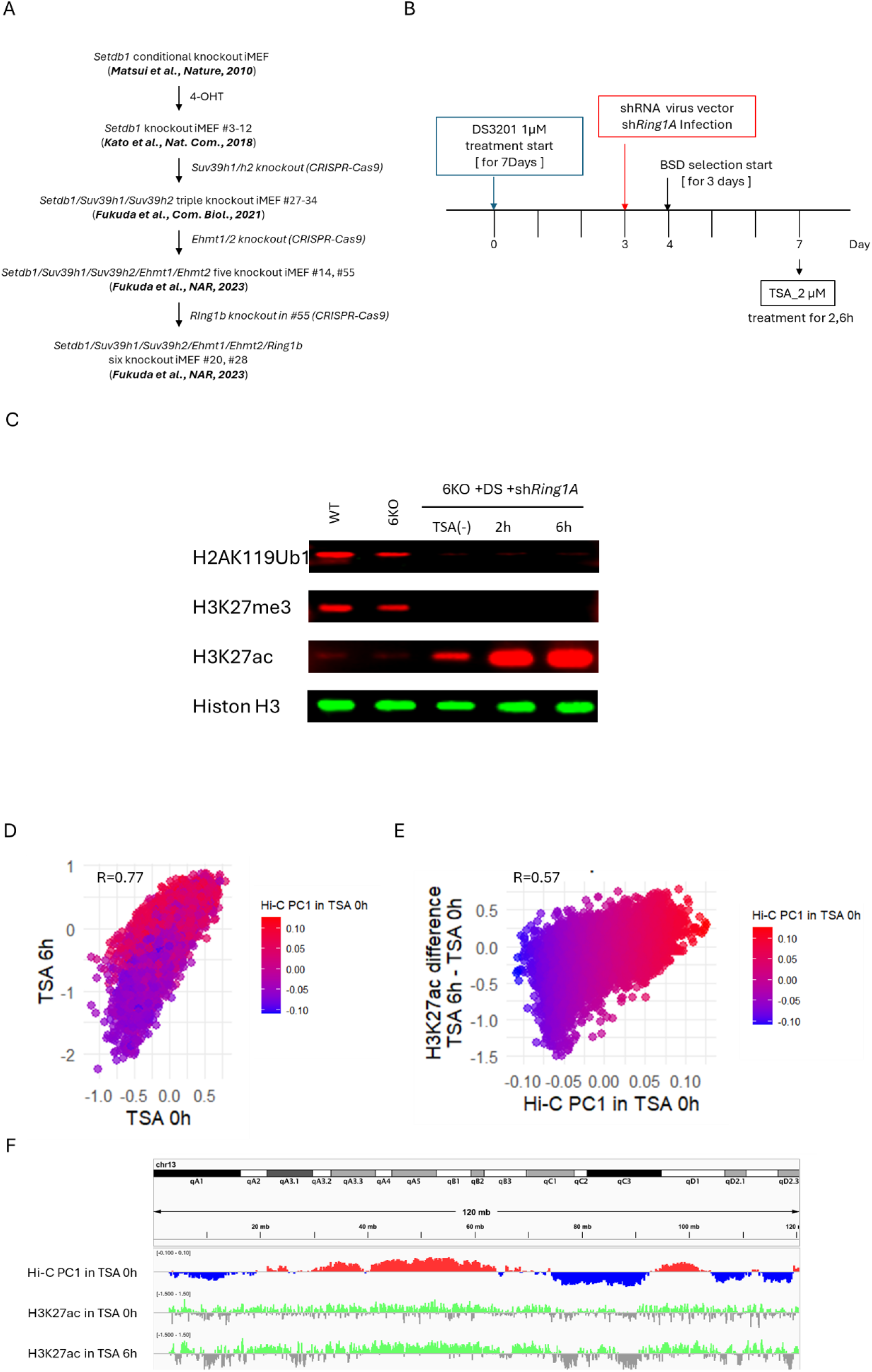
Differences in H3K27ac increase between compartments induced by TSA treatment. (A) Scheme of 6KO iMEFs establishment. (B) Scheme of DS3201 treatment, Ring1a knockdown, and TSA treatment on 6KO iMEFs. (C) Western blot analysis of H3K27ac, uH2A and H3K27me3 levels. (D) Scatter plot of H3K27ac enrichment. The log2(ChIP/Input) values in 250-kb bins were compared between TSA 0 h and 6 h. Dot colors indicate the Hi-C PC1 value at TSA 0 h: red represents the A compartment, and blue represents the B compartment. (E) Scatter plot of Hi-C PC1 values and difference in H3K27ac between samples. X-axis represents Hi-C PC1 values in TSA 0h and Y-axis represents the difference in log2(ChIP/Input) of H3K27ac between TSA 0 h and 6 h for each 250-kb bin. Dot colors indicate the Hi-C PC1 value at TSA 0 h: red represents the A compartment, and blue represents the B compartment. (F) Representative genomic regions illustrating the relationship between the H3K27ac profile and Hi-C PC1 value.

Histone Deacetylases(HDACs)are also important factors for heterochromatin assembly (Allshire and Madhani 2018; Fukuda and Shinkai 2020; Bell et al. 2023; Grewal 2023). TSA is a potent HDAC inhibitor (Yoshida et al. 1990), and treatment of mammalian cells with TSA induces chromocenter disassembly and upregulation of transposon expression that is normally repressed by H3K9 methylation (Taddei et al. 2001; Kato et al. 2018). Therefore, we treated 6KO cells, in which H3K27 methyltransferase inhibitor DS3201 and *Ring1a* were knocked down (6KO+sh1A+DS), with TSA and examined its effects (Fig. 1B). Treatment with 2 µM TSA for 2 and 6 hours significantly increased H3K27ac levels (Fig. 1C). In addition, H3K27me3 and uH2A levels were markedly decreased, confirming that simultaneous treatment with TSA, *shRing1A* and DS3201 achieved both H3K27me3/uH2A reduction and H3K27ac increase (Fig. 1C). Since prolonged TSA treatment beyond 6 hours induced cell death, we decided to continue the analysis under short term TSA treatment conditions.

To investigate the genome-wide profile of H3K27ac and 3D genome architecture following TSA treatment, we performed H3K27ac ChIP-seq and Hi-C on 6KO+sh1A+DS cells treated with TSA for 0 or 6 hours. When comparing the log_2_(ChIP/Input) values of 250-kb bins between 0 h and 6 h, a positive correlation was observed (R=0.77), indicating that the state of high H3K27ac in A compartments and low H3K27ac in B compartments was maintained (Fig. 1D). Additionally, TSA treatment tended to increase H3K27ac levels specifically in A compartments (R=0.57) (Fig. 1E, F), implicating A compartment preference of TSA.

Chromocenter analysis using DAPI staining revealed no reduction in Chromocenter formation upon TSA treatment (Fig. 2A). Furthermore, Hi-C analysis showed no significant changes in the Pearson correlation matrix or interaction matrix after TSA treatment (Fig. 2B, C). The Hi-C PC1 values were also unaffected by TSA treatment (R = 0.98) (Fig. 2D). A-to-B or B-to-A compartment conversions remained below 4% following TSA treatment (Fig. 2E), indicating that the increase in H3K27ac induced by TSA did not result in substantial changes in nuclear compartments.

**Figure 2.**
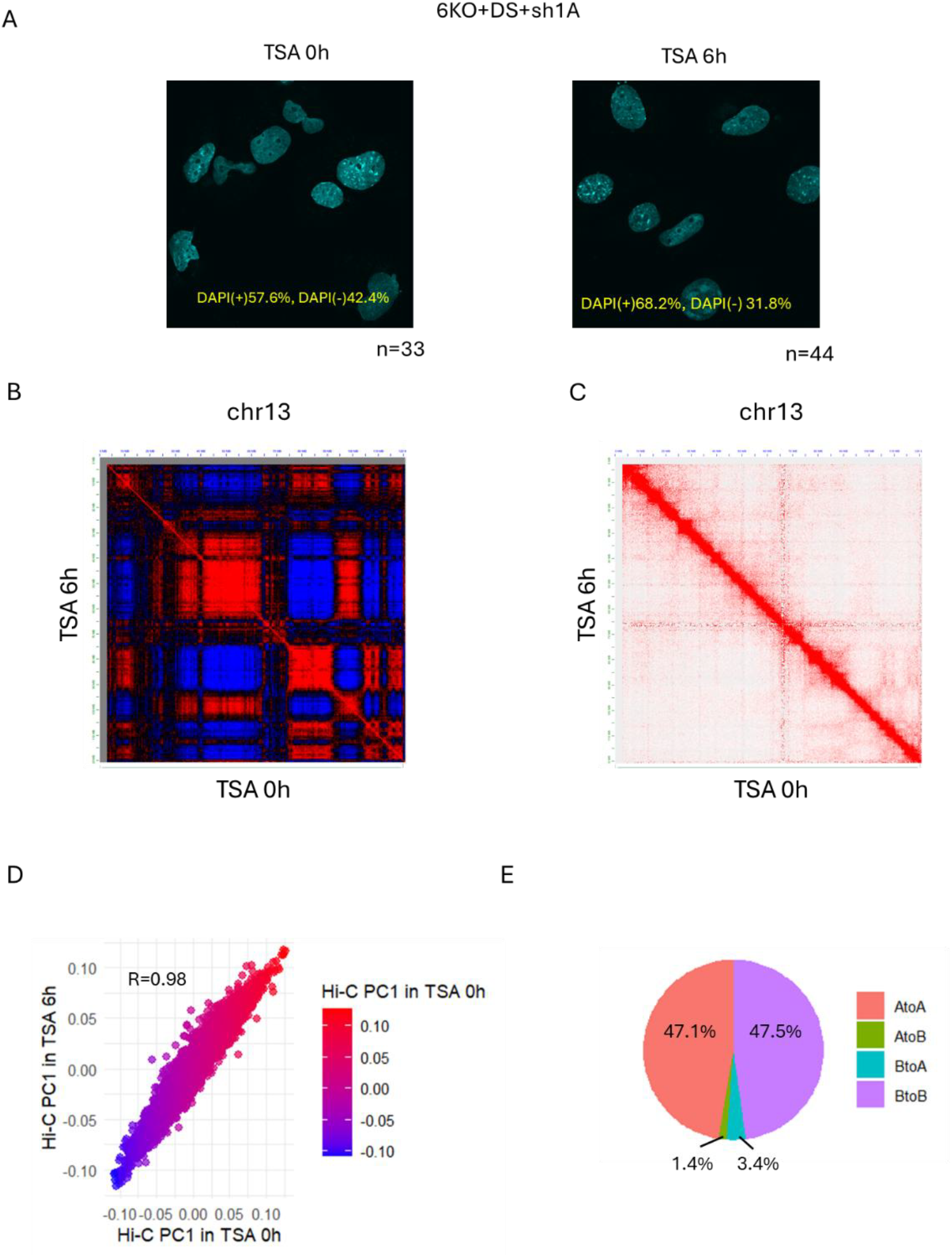
Prevention of abnormal CTCF bindings at repetitive elements by H3K9/K27 methylation. (A) DAPI-stained cell images. Cells with and without DAPI-dense foci (chromocenters) were counted. (B, C) Pearson correlation matrix (B) and interaction matrix (C) on chromosome 13. (D) Scatter plot of Hi-C PC1 values between 0h and 6h. Dot colors indicate the Hi-C PC1 value at TSA 0 h: red represents the A compartment, and blue represents the B compartment. Hi-C PC1 values of 6h is highly correlated with those of 0h (Pearson’s R = 0.98). (E) Pie chart of compartment conversion after 6h treatment of TSA.

Our study demonstrated that the increase in H3K27ac induced by TSA treatment does not significantly affect nuclear compartments in cells depleted of H3K9 methylation, H3K27me3, and uH2A. The increase in H3K27ac following TSA treatment mainly occurred within A compartments, while the quantitative difference in H3K27ac levels between A and B compartments was maintained. This may partly explain the limited impact on nuclear compartmentalization.

Previous studies have reported that chromocenter disassembly following TSA treatment occurs after prolonged exposure, such as 5 days (Taddei et al. 2001). In 6KO+sh1A+DS condition, achieving increased H3K27ac in B compartments or significant alterations in nuclear compartments may also require long-term TSA treatment. However, extended TSA exposure in this condition induces cell death, making it challenging to investigate. Why does TSA preferentially increase H3K27ac in A compartments? Is it due to differences in HDAC or HAT activity or abundance between compartments, or is transcriptional activity the key factor? Elucidating these underlying mechanisms could lead to a deeper understanding of compartment-specific differences.

## MATERIALS AND METHODS

### Cell culture

We used previously established *Setdb1, Suv39h1/2, Ehmt1, Ehmt2, Ring1b* KO iMEFs to analysis the role of uH2A in heterochromatin maintenance (Fukuda et al. 2021). Mouse embryonic fibroblasts were maintained in Dulbecco’s modified Eagle’s medium (Nacalai tesque, 08458-16) containing 10% fetal bovine serum (Biosera, FB1061), MEM Non-Essential medium and 2-Mercaptoethanol (Nacalai tesque, 21417-52). To inhibit EZH1/2 catalytic activity, iMEFs were cultured for seven days with 1 *μ*M DS3201.

### *Ring1a* knockdown by shRNA

We produced a lentiviral vector expressing shRNA targeting *Ring1a* by transfecting 293FT cells with shRNA vector, psPax2, and pMD2.G using PEI. Two days later, we collected the culture supernatant and then used it to transduce 5KO-*Ring1b* KO iMEFs at MOI=2. Following selection with 7 μg/ml BSD for 3 days, we collected the cells.

### TSA treatment

Six knockout (6KO) iMEFs treated with DS3201 and shRing1A were exposed to 2 μM TSA for 2 hours and 6 hours, followed by Western blotting, H3K27ac ChIP-seq, and in situ Hi-C analysis.

### H3K27ac ChIP-seq

Native ChIP and crosslinked ChIP. Native ChIP assays were performed as described previously (Fukuda et al. 2023). Antibody against H3K27ac (D5E5, CST) was used. The ChIP DNA was fragmented by Picoruptor (Diagenode) for 10 cycles of 30 seconds on, 30 seconds off. Then, ChIP library was constructed by KAPA Hyper Prep Kit (KAPA BIOSYSTEMS) and SeqCap Adapter Kit A (Roche) according to manufacturer instructions. The concentration of the ChIP-seq library was quantified by KAPA Library quantification kit (KAPA BIOSYSTEMS). ChIP sequencing was performed on a HiSeq X platform (Illumina). We performed two biological replicates for ChIP-seq.

### ChIP-seq analysis

Adaptor sequences and low quality bases in reads were trimmed using Trim Galore version 0.3.7 (http://www.bioinformatics.babraham.ac.uk/projects/trim_galore/). Then trimmed reads were aligned to the mouse GRCm38 genome assembly using bowtie version 0.12.7 (Langmead 2010) with default parameters. Duplicated reads were removed using samtools version 0.1.18 (Li et al. 2009).

### Preparation of Hi-C library

Hi-C experiments were performed as previously described (Ikeda et al. 2018; Kadota et al. 2020), based on DpnII enzyme (4-bps cutter) using 2×10^6^ fixed cells. Hi-C libraries were subject to paired-end sequencing (150 base pair (bp) read length) using HiSeq X Ten. Detailed protocol for HiC-seq library preparation is available at Protocols.io (https://www.protocols.io/view/iconhi-c-protocol-ver-1-0-4mjgu4n).

### Hi-C data analysis

Hi-C data processing was done by using Docker for 4DN Hi-C pipeline (v43, https://github.com/4dn-dcic/docker-4dn-hic). The pipeline includes alignment (using the mouse genome, mm10) and filtering steps. After filtering valid Hi-C alignments, .*hic* format Hi-C matrix files were generated by Juicer Tools (Durand et al. 2016) using the reads with MAPQ>10. The A/B compartment (compartment score) profiles (in 250 kb bins) in each chromosome (without sex chromosome) were calculated from .*hic* format Hi-C matrix files (intrachromosomal KR normalized Hi-C maps) by Juicer Tools (Durand et al. 2016) as previously described (Miura et al. 2018). We averaged Hi-C PC1 values in each 250 kb bin from two biological replicates for the downstream analysis.

### Visualization of NGS data

The Integrative Genomics Viewer (IGV) (Robinson et al. 2011) was used to visualize NGS data. For Hi-C contact matrix and correlation matrix, we used Juicer Tools (Durand et al. 2016).

## Data availability

All NGS data used in this study have been deposited in the Gene Expression Omnibus (GEO) under accession number GSE294421 and GSE294422.

## ACKNOWLEDGEMENTS

We thank the staff of the Support Unit for Bio-Material Analysis (BMA) at the RIKEN Center for Brain Science (CBS) Research Resources Division (RRD) for NGS library construction, DNA sequencing and flow cytometry. We would also like to thank our colleagues at Shinkai laboratory for their support and valuable comments.

## Author contributions

K.F. and Y.S. designed and conceived the study. K.F. and Y.S. supervised the study and interpreted the data. C.S. performed molecular and cellular experiments and generated the ChIP-seq, RNA-seq and Hi-C-seq libraries. K.F. performed informatics analysis of generated NGS data. K.F. and Y.S. wrote the manuscript and prepared figures. All authors read, discussed, and approved the manuscript.

